# Integrative multi-omic analysis identifies genes associated with cuticular wax biogenesis in adult maize leaves

**DOI:** 10.1101/2024.04.09.588685

**Authors:** Meng Lin, Harel Bacher, Richard Bourgault, Pengfei Qiao, Susanne Matschi, Miguel F. Vasquez, Marc Mohammadi, Sarah van Boerdonk, Michael J. Scanlon, Laurie G. Smith, Isabel Molina, Michael A. Gore

**Affiliations:** Plant Breeding and Genetics Section, School of Integrative Plant Science, Cornell University, Ithaca, NY 14853, USA; Department of Biology, Algoma University, Sault Ste. Marie, ON P6A 2G4, Canada; Plant Biology Section, School of Integrative Plant Science, Cornell University, Ithaca, NY 14853, USA; Department of Cell and Developmental Biology, University of California San Diego, La Jolla, CA 92093, USA

**Keywords:** Maize, Leaf cuticular waxes, Leaf cuticular conductance, Natural variation, Genome-wide association study, Transcriptome-wide association study, Fisher’s combined test, Random forest, Genetic architecture, GWAS hotspot

## Abstract

Studying the genetic basis of leaf wax composition and its correlation with leaf cuticular conductance (*g*_c_) is crucial for improving crop water-use efficiency. The leaf cuticle, which comprises a cutin matrix and various waxes, functions as an extracellular hydrophobic layer, protecting against water loss upon stomatal closure. To address the limited understanding of genes associated with the natural variation of leaf cuticular waxes and their connection to *g*_c_, we conducted statistical genetic analyses using leaf transcriptomic, metabolomic, and physiological data sets collected from a maize (*Zea mays* L.) panel of ∼300 inbred lines. Through a random forest analysis with 60 cuticular wax traits, it was shown that high molecular weight wax esters play an important role in predicting *g*_c_. Integrating results from genome-wide and transcriptome-wide studies (GWAS and TWAS) via a Fisher’s combined test revealed 231 candidate genes detected by all three association tests. Among these, 11 genes exhibit known or predicted roles in cuticle-related processes. Throughout the genome, multiple hotspots consisting of GWAS signals for several traits from one or more wax classes were discovered, identifying four additional plausible candidate genes and providing insights into the genetic basis of correlated wax traits. Establishing a partially shared genetic architecture, we identified 35 genes for both *g*_c_ and at least one wax trait, with four considered plausible candidates. Our study uncovered the genetic control of maize leaf waxes, establishing a link between wax composition and *g*_c_, with implications for potentially breeding more water-use efficient maize.

**SIGNIFICANCE STATEMENT:** We exploited natural variation in the abundance of maize leaf cuticular waxes to identify genetic determinants of wax composition and its relationship to cuticle function as a barrier against water loss. We identified a set of strongly supported candidate genes with plausible functions in cuticular wax biosynthesis or deposition and added to the evidence for wax esters as the most important wax for water barrier function, offering new tools for modification of cuticle-dependent traits.

## INTRODUCTION

The cuticle covers the surface of aerial organs of all land plants. Being predominantly hydrophobic, cuticles provide a near-complete barrier from the environment, protecting internal tissues from biotic and abiotic stresses (Bargel *et al*., 2004). For instance, the cuticle can reduce pathogen (Serrano *et al*., 2014) and insect susceptibility (Eigenbrode, 2002), radiation damage (Krauss *et al*., 1997), and limit water loss (Kerstiens, 1996; Riederer and Schreiber, 2001). Approximately 5-10% of water lost by well-hydrated, transpiring plants during daylight hours is estimated to be attributed to evaporation occurring through the cuticle (Taiz and Zeiger, 2010). However, essentially all water lost at night, and under water limiting conditions where stomata are closed, is lost via cuticular evaporation, or through stomatal pores that are not completely sealed (Schuster *et al*., 2017; Resco de Dios *et al*., 2019).

Leaf cuticles consist of mixtures of waxes and cutin, although their structure and composition vary between and within species and developmental stages (Buschhaus *et al*., 2007; Fernández *et al*., 2016; Pollard *et al*., 2008; Goodwin and Jenks, 2005). Cuticular waxes and cutin are organized into three layers. The deepest layer, which is continuous with the cellulosic wall, is called the cuticular layer, consisting of polysaccharides and cutin. The middle layer, or cuticle proper, is a highly hydrophobic matrix of cutin embedded with intracuticular waxes. The outermost layer is a film of epicuticular waxes (Domínguez *et al*., 2011; Albersheim *et al*., 2010). Several studies have shown that waxes are the main cuticle component that prevents non-stomatal water loss (Leide *et al*., 2007; Schönherr and Riederer, 1989; Bi *et al*., 2017), serving as a barrier to water diffusion across the leaf (Seufert *et al*., 2021).

Cuticular waxes consist of solvent-extractable compounds including very long-chain fatty acids (FAs), alcohols (including primary alcohols, PAs), hydrocarbons (HCs), aldehydes (ADs), wax esters (WEs), and alicyclics (ACs) (Jetter *et al*., 2018). In contrast, cutin is an insoluble polymer mainly formed by ester-bonded C16 and C18 hydroxy and hydroxy-epoxy fatty acid monomers (Kolattukudy, 1970). Interspecies comparisons and mutant characterization have provided evidence that leaf water conductance is neither determined by the total amount of wax nor the thickness of the cuticle, but rather by the relative proportions and organization of the waxes (Riederer and Schreiber, 2001). Cuticular waxes are formed in the endoplasmic reticulum (ER), where CoA-thioesters of plastidial fatty acids are elongated to very long-chain acyl-CoAs by the multi-enzyme fatty acid elongase complex (Jenks *et al*., 1995). Very long-chain acyl-CoAs are either hydrolyzed to become long-chain free fatty acids (Lü *et al*., 2009), or may enter one of two pathways: an acyl reduction pathway that produces primary alcohols and wax esters, or an alkane-forming pathway that produces aldehydes, alkanes, secondary alcohols, and ketones (Yeats and Rose, 2013). Waxes are delivered to the plasma membrane via vesicular transit through the Golgi (McFarlane *et al*., 2014) and across the plasma membrane by ABC transporters (McFarlane *et al*., 2010). The transport of waxes across the plasma membrane is also facilitated by glycosylphosphatidylinositol-anchored lipid-transfer proteins (DeBono *et al*., 2009; Lee *et al*., 2009; Kim *et al*., 2012).

The underlying genetics of cuticular wax biogenesis and transport has been initially advanced by studying plant mutants. For example, the characterization of “glossy mutants” in grasses, including sorghum [*Sorghum bicolor* (L.) Moench], barley (*Hordeum vulgare* L.), and maize (*Zea mays* L.), has revealed some genes that are critical for wax biosynthesis and deposition (Jenks *et al*., 1994; Lundqvist and Lundqvist, 2008; Schnable *et al*., 1994; Bianchi *et al*., 1979). In maize, more than 30 cuticular wax mutants have been identified, showing a glossy leaf phenotype (Schnable *et al*., 1994); however, only some of these genes have been cloned and characterized. Genes associated with glossy phenotypes, including *GLOSSY1* (*GL1*), *GL2*, *GL4*, *GL8*, and *GL26*, encode enzymes predicted to be involved in cuticular wax biosynthesis (Sturaro *et al*., 2005; Tacke *et al*., 1995; Liu *et al*., 2009; Dietrich *et al*., 2005; Xu *et al*., 2002; Fan, 2007). Furthermore, *GL3* and *GL15* encode MYB and AP2-like transcription factors, respectively, regulating cuticular wax biosynthesis (Moose and Sisco, 1994; Liu *et al*., 2012). Contrastingly, *GL6* and *GL13* encode the DUF538 protein and an ABC transporter, respectively, involved in wax transport (Li *et al*., 2019; Li *et al*., 2013), while *GL14* encodes a putative membrane-associated protein with a yet-to-be-discerned function (Zheng *et al*., 2019).

While many genes involved in the biosynthesis, transport, and deposition of waxes have been reported, studies at the genome-wide level have been limited to investigating the genetic control of natural variation for the abundances of cuticular waxes in camelina [*Camelina sativa* (L.) Crantz] and sorghum (Z., Luo *et al*., 2019; Elango *et al*., 2020; Jin *et al*., 2020). Our prior studies in maize have primarily focused on identifying candidate genes associated with cuticle-dependent water loss (Lin *et al*., 2022; Lin *et al*., 2020), a phenomenon we term adult leaf cuticular conductance (*g*_c_) (Lin *et al*., 2020). By integrating findings from both genome- and transcriptome-wide association studies (GWAS and TWAS) via the Fisher’s combined test (FCT) in a panel of 310 maize inbred lines, Lin *et al*. (2022) revealed the association of *g*_c_ with 22 plausible candidate genes, including those predicted to play roles in cuticle precursor biosynthesis and export, intracellular membrane trafficking, and the regulation of cuticle development. With a subset of ∼50 lines having the highest and lowest *g*_c_ scores, Lin *et al*. (2022) showed that leaf cuticular wax composition of these lines is moderately predictive of their *g*_c_ values, implying that levels of specific cuticular waxes affect cuticle permeability.

To enrich the depth of our initial efforts, we combined GWAS and TWAS results to thoroughly analyze natural variation in the abundance of 60 leaf cuticular wax traits in the complete panel of 310 maize inbred lines that Lin *et al*. (2022) scored for *g*_c_. Our specific aims were to: (*i*) determine the relationship of cuticular waxes to *g*_c_, (*ii*) identify candidate genes associated with wax traits, (*iii*) detect genomic hotspots that control multiple wax traits, and (*iv*) reveal candidate genes shared between waxes and *g*_c_.

## RESULTS

### Phenotypic analysis

The abundance of 53 cuticular wax compounds (7 PAs; 11 FAs; 9 HCs; 4 ADs; 14 WEs; and 8 ACs) and seven summed traits (total mass per unit area of compounds in each of the six classes and the total mass of all compounds per unit area) were assessed across 310 genetically diverse maize inbred lines of the Wisconsin Diversity (WiDiv) panel (Table S1). The measured cuticular wax components showed a wide range of abundances, with average concentrations ranging from 0.07 (HC 39:0) to 13.25 (HC 31:0) μg·dm^-2^. Of the six class sum traits, total HC was the most abundant (39.12 μg·dm^-2^), whereas total AD was the least abundant (3.63 μg·dm^-2^). However, total AC had a coefficient of variation (CV) of 0.44, which was the highest CV observed among the six sum traits. The 60 cuticular wax traits had an average heritability of 0.70, ranging from 0.17 (FA 18:0) to 0.97 (AC Friedelin, AC Unk1, and AC Unk7) (Table S1), indicating that variation for cuticular wax abundance among lines is predominantly under genetic control. On average, the pairwise Pearson’s correlations for the 53 cuticular wax compounds revealed moderately strong correlations (average *r* = 0.34) between compounds belonging to the same class. In contrast, correlations were relatively weaker (average *r* = 0.12) between compounds belonging to different classes (Figure S1; Table S2). Altogether, the stronger correlations observed between compounds of the same class suggest a shared genetic architecture, possibly indicating a common biosynthetic pathway.

### Prediction of *g*_c_ with cuticular waxes

Lin *et al*. (2022) assessed cuticular conductance (*g*_c_), a measure of water loss resulting from leaf cuticle permeability, in the same collection of leaves whose wax profiles are reported here. Recognizing that the content and composition of leaf cuticular waxes may impact cuticle permeability, we investigated the relationship between *g*_c_ to the 60 cuticular wax traits. We found significant Pearson’s correlations of 18 cuticular wax traits (including total wax and a subset of FAs, HCs, WEs and ACs) with *g*_c_ (|*r*| = 0.13 - 0.24; *P*-value < 0.05). Among these 18 wax traits, only HC 27:0 showed a positive correlation with *g*_c_ (*r* = 0.14; Figure S1; Table S2). Interestingly, all 10 significant WEs had a negative correlation with *g*_c_ (Table S2), suggesting that a higher abundance of these WEs could potentially reduce the permeability of the leaf cuticle to water.

To provide a more comprehensive assessment of the relationship between *g*_c_ and leaf cuticular waxes, a RF analysis was performed to assess the collective ability of wax traits to predict *g*_c_ and determine their individual importance in the prediction model. The 60 cuticular wax traits, when combined, demonstrated an overall predictive ability of 0.27 for *g*_c_. Comprising predominantly high molecular weight WEs, only 10% of the top-ranked features (in descending order of importance: WE 54:0, FA 22:0, WEs 49:0, 50:0, 52:0, 47:0) contributed the majority of the predictive power, as indicated by their importance scores (Figure 1). Suggesting stability in the RF analysis, we observed a robust positive correlation (*r* = 0.48; *P*-value = 1.03 × 10^-4^) between the ranks of importance scores for cuticular waxes in this analysis and a comparable analysis undertaken by Lin *et al*. (2022) using a subset of 51 WiDiv lines with extremely high or low *g*_c_ values (Table S3). Further underscoring the importance of cuticular wax composition rather than overall wax amount in predicting *g*_c_, we observed a non-significant, weak negative correlation between the importance scores and the average abundances of leaf cuticular waxes (*r* = -0.17; *P*-value = 0.22). This implies that the relative abundance of different waxes did not appear to significantly impact their importance in predicting *g*_c_. While waxes collectively account for only a moderate amount of the total variation in *g*_c_, a genetic dissection of wax composition would contribute to a deeper understanding of their genetic regulation and their role in influencing *g*_c_ variability.

**Figure 1.**
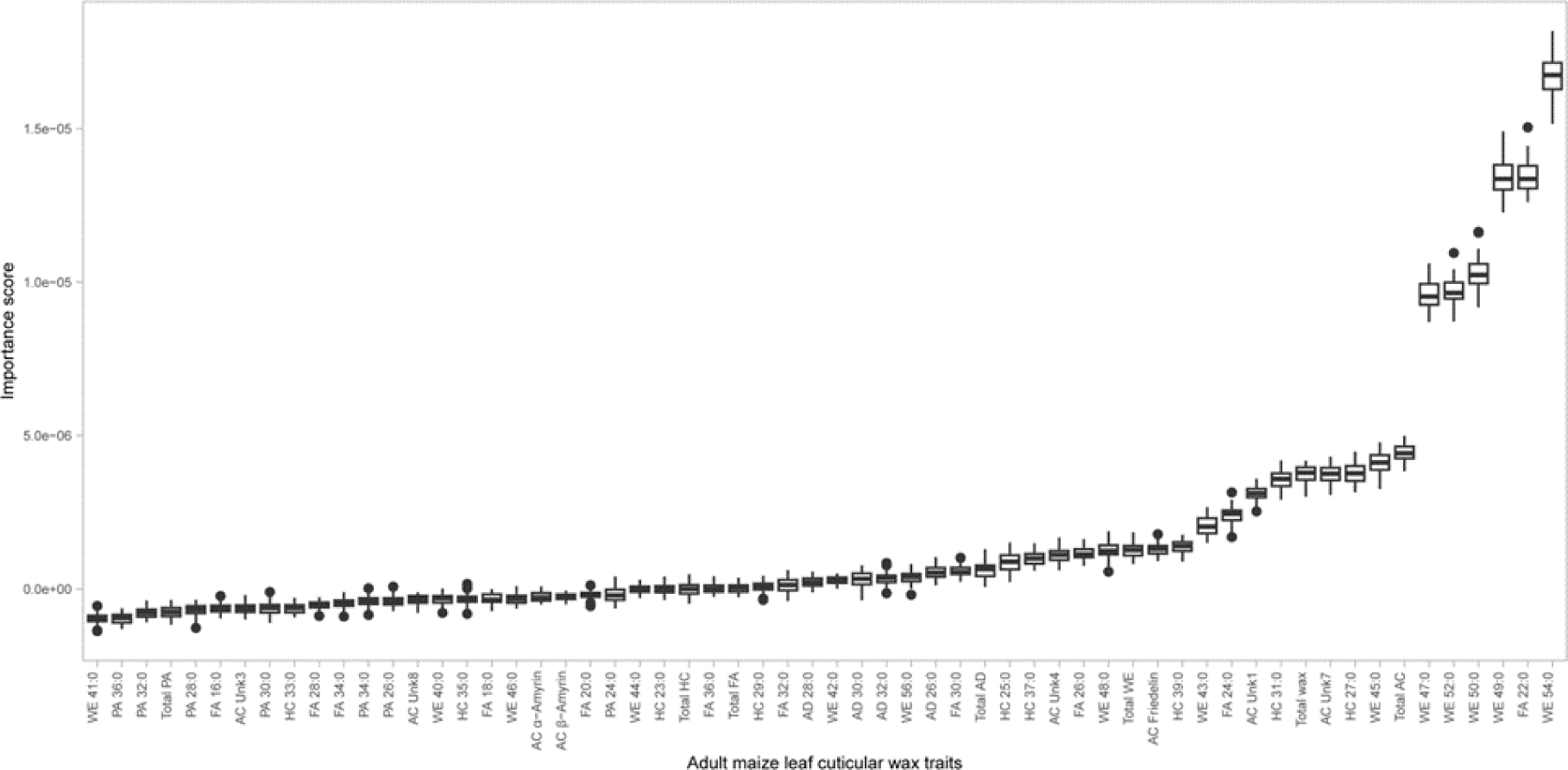
Box plots of importance scores for cuticular wax composition in random forest regression to predict maize leaf cuticular conductance (*g*_c_). Box limits indicate the upper and lower quartiles; center lines in boxes indicate the median value; whiskers indicate 1.5x interquartile range; and dots indicate outliers.

### Quantitative genetic analysis for identification of candidate genes for cuticular waxes

We performed GWAS and TWAS on the 60 cuticular wax traits to investigate the genetic basis of cuticular wax accumulation in the maize adult leaf, and then integrated the GWAS and TWAS results with the FCT to enhance statistical power. A total of 6,186 unique candidate genes (top 0.002% SNPs) were identified from GWAS (Tables S4 and S5), 1,956 (top 0.25% genes) from TWAS (Table S6) and 2,965 (top 0.25% genes) from FCT (Table S7) across the 60 wax traits. The number of unique candidate genes identified across the three methods varied from 103 (AC α-Amyrin & AC β-Amyrin) to 398 (AD 30:0) for each of the 60 traits (Table S8). Of the identified candidate genes, 231 were designated as “higher confidence” candidates (Table S9) because they were detected by FCT, TWAS, and GWAS when collectively considering all 60 wax traits. Of these, 69 (∼30%) were associated with one or more of the six wax compounds that were ranked by the RF analysis as being of the highest importance in predicting *g*_c_ (in descending order of importance: WE 54:0, FA 22:0, WEs 49:0, 50:0, 52:0, 47:0). As shown in Table S9, the higher confidence candidate genes were divided into three groups using the following criteria: group 1) consisted of 41 genes identified by FCT, TWAS, and GWAS for the same trait; group 2) comprised 42 genes not meeting criteria of group 1 but identified by FCT, TWAS, and GWAS when collectively considering all traits within a wax class; and group 3) included 148 genes not meeting the criteria for groups 1 and 2 but identified by all 3 methods. Of the 231 higher confidence candidates listed in Table S9, 11 genes are featured as plausible candidates in Table 1 based on predicted functions that can be related to cuticular waxes. Five of these (*Zm00001d045660*, *Zm00001d018474*, *Zm00001d038404/Zm00001d038405*, *Zm00001d013798*, and *Zm00001d049100*) were associated with at least one of the six compounds found to be most important for the *g*_c_ trait via RF analysis. The closest Arabidopsis [*Arabidopsis thaliana* (L.) Heynh.] and rice (*Oryza sativa* L.) homologs of these 11 genes along with additional details are listed in Table S10.

**Table 1.**
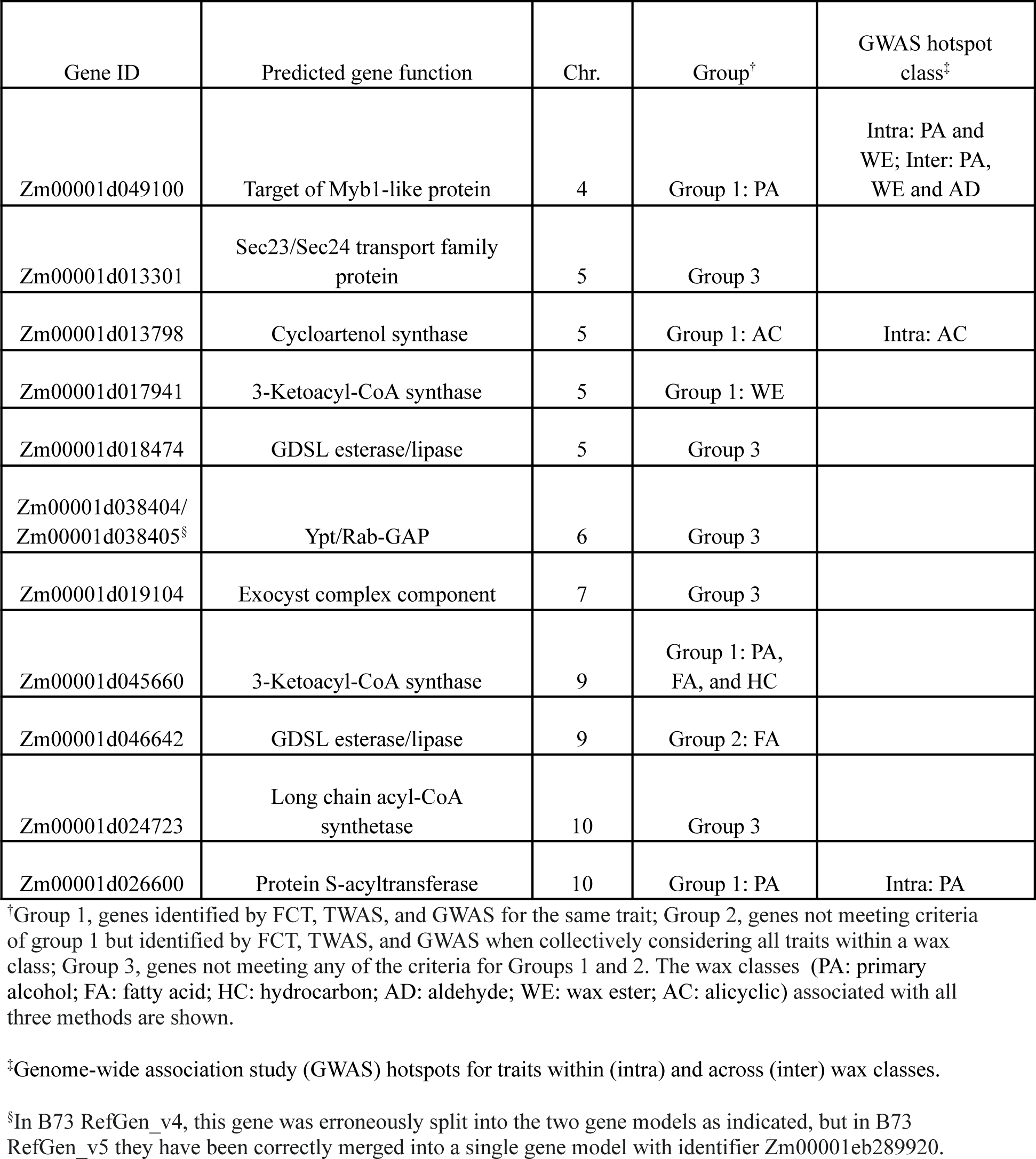
Plausible candidate genes identified for adult maize leaf cuticular waxes via a genome-wide association study (GWAS), transcriptome-wide association study (TWAS), Fisher’s combined test (FCT), and GWAS hotspot analysis in the maize Wisconsin diversity panel. An expanded version of this table with additional information about each gene is provided as Table S10.

Of the higher confidence candidate genes, six listed in Table 1 are predicted or functionally confirmed in maize to have a role in cuticular wax biosynthesis (Table S10). Two members of the 3-ketoacyl-CoA synthase (KCS) family, namely KCS22 encoded by *Zm00001d045660* (*KCS22*) and a KCS6 paralog encoded by *Zm00001d017941* (*GLOSSY4B*, *GL4B*), were identified in association with 26 and six unique wax traits, respectively. In TWAS and FCT, *KCS22* exhibited associations with four (WEs 49:0, 50:0, 52:0, 54:0; Figure 2) of the six compounds identified as most important for *g*_c_ through RF analysis. However, no significant associations with these six compounds were observed for *GL4B*. *Zm00001d013798*, which encodes a homolog of Arabidopsis CYCLOARTENOL SYNTHASE1 (CAS1) (Corey *et al*., 1993; Babiychuk *et al*., 2008), was identified in GWAS for five ACs and 11 unique traits (including two FAs and nine ACs) when combining TWAS and FCT results. Two additional higher confidence candidate genes listed in Table 1 encoding putative Gly-Asp-Ser-Leu (GDSL) lipases, are each associated with three unique wax traits. One of these, *Zm00001d046642* (encodes a homolog of Arabidopsis OCCLUSION OF STOMATAL PORE1, OSP1), showed associations with FA 24:0 in TWAS, and with FA 36:0 and HC 35:0 in both GWAS and FCT. The other, *Zm00001d018474* (*GDSL ESTERASE/LIPASE2*, *GELP2*) exhibited associations with HC 27:0 in TWAS and FCT, as well as with WEs 49:0 and 56:0 in GWAS (Figure 2). Finally, the *CER8* gene (*Zm00001d024723*), encoding a homolog of Arabidopsis CER8/LONG-CHAIN ACYL-COA SYNTHETASE1 (LACS1) (Lü *et al*., 2009), was associated with PA 32:0 in GWAS, and with HC 37:0 in both TWAS and FCT.

**Figure 2.**
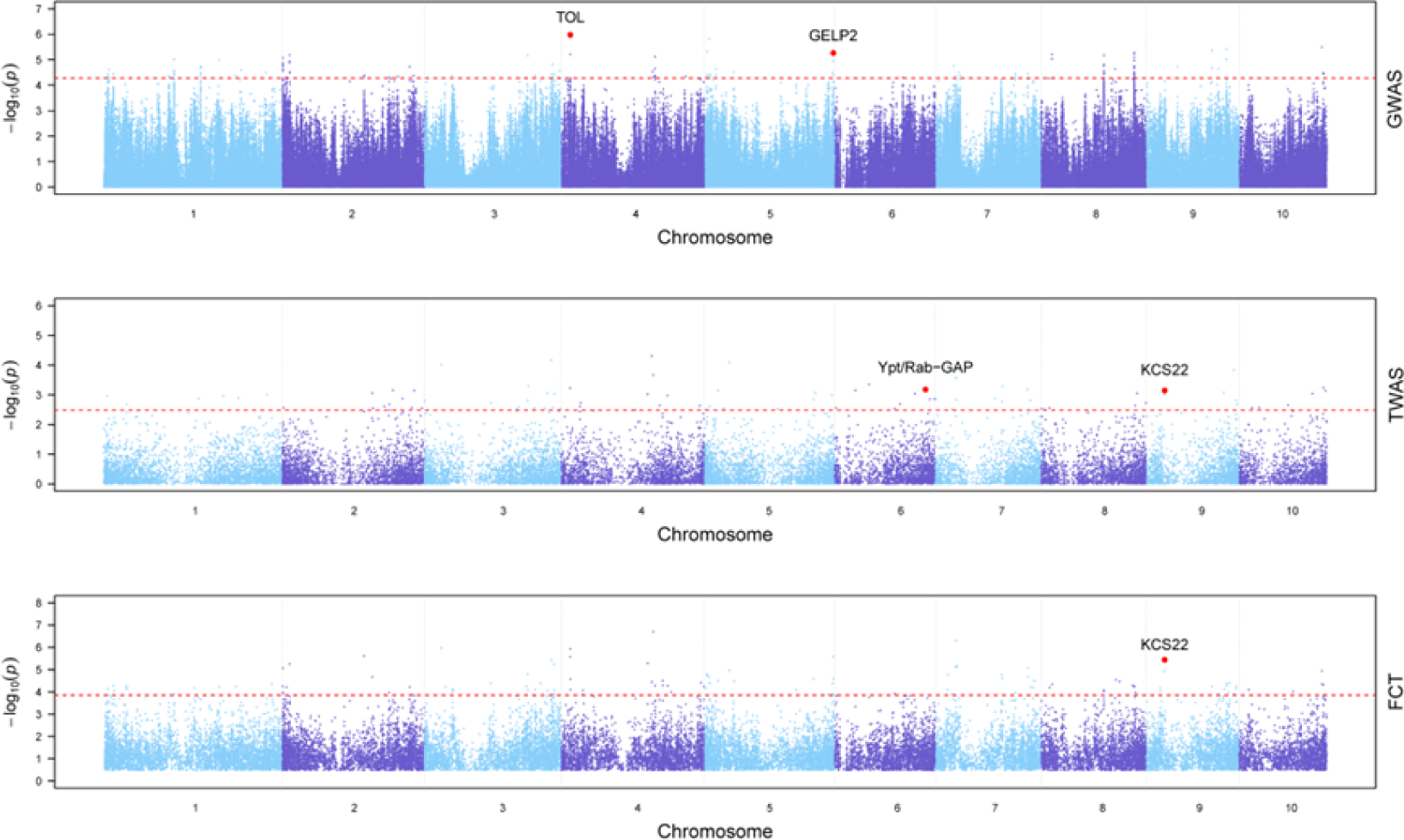
Manhattan plots of GWAS, TWAS, and FCT results for WE 49:0. Each point represents a SNP or gene with its −log_10_ *P*-value (y-axis) from GWAS, TWAS, and FCT plotted as a function of physical position (Mb, B73 RefGen_v4) across the 10 chromosomes of maize (x-axis). Red horizontal dashed lines indicate the thresholds of top 0.002%, top 0.25%, and top 0.25% for GWAS, TWAS, and FCT, respectively. Plausible candidate genes (Table 1) that are within 200 kb of a top 0.002% GWAS peak SNP or ranked in the top 0.25% in TWAS or FCT are highlighted with red dots and labeled in black in the Manhattan plots.

The transport of waxes from their synthesis site in intracellular membranes to the cell wall surface depends on Golgi-mediated vesicular trafficking (McFarlane *et al*., 2014). Accordingly, the remaining five of 11 higher confidence candidate genes listed in Table 1 were chosen because they encode proteins predicted to function in intracellular trafficking as discussed later. *Zm00001d038404*/*Zm00001d038405*, which encodes a protein with a putative Ypt/Rab-GTPase-activating protein (GAP) domain, was found to be associated with nine unique traits. Notably, among these traits, four (WEs 49:0, 50:0, 52:0, 54:0; Figure 2) were found to be of high importance in the RF analysis. *Zm00001d019104*, encoding a homolog of Arabidopsis exocyst complex component 84B (EXO84B), was associated with FA 32:0 and HC 39:0. *Zm00001d013301* (*SALT-TOLERANCE-ASSOCIATED-GENE4*, *SAG4*) (X., Luo *et al*., 2019), which associated with FA 36:0 and three HC (33:0, 35:0, and total HC) traits, encodes a SECRETION23/SECRETION24 (SEC23/SEC24) family protein. *Zm00001d026600* (*PROTEIN S-ACYLTRANSFERASE38*, *PAT38*), which encodes a protein with similarity to members of the DHHC-type zinc finger protein family in Arabidopsis that mediate the S-acylation of proteins (Yuan *et al*., 2013), was found to be associated with eight distinct traits (six PAs, AD 28:0, and WE 48:0). *Zm00001d049100*, encoding a protein similar to members of the Arabidopsis TARGET OF MYB1-LIKE (TOL) family of proteins involved in the sorting of ubiquitinated protein cargoes into multivesicular bodies (Winter and Hauser, 2006), was associated with 11 AD, PA and WE traits (Figure 2).

Identification of additional candidate genes was facilitated by cross-referencing our GWAS, TWAS, and FCT results against a list of 115 candidate genes for maize leaf cuticle biosynthesis and its regulation, as compiled by Qiao *et al*. (2020) based on transcriptome analysis. This resulted in the identification of 39 candidate genes from the list of Qiao *et al*. (2020) that were associated with at least one wax trait (Table S11). Four of these genes (*Zm00001d018474, GELP2*; Zm00001d024723, *CER8*; *Zm00001d045660, KCS22*; *Zm00001d046642*, *OSP1* homolog) are included in our list of higher confidence candidate genes, as they were detected by GWAS, TWAS, and FCT (Table S9). The remaining 35 candidate genes, identified by two or fewer of the three methods, were collectively found to be associated with a total of 32 traits from all six wax classes. Although these 35 candidate genes have less robust support compared to those detected by all three methods, they are still well-supported as candidates having a purported role in the genetic control of wax traits. In the context of the six top-ranked wax compounds revealed by the RF analysis as having high importance for predicting *g*_c_, five of the 35 candidate genes (*GLYCEROL-3-PHOSPHATE ACYLTRANSFERASE20*, *GPAT20*; *KCS24*; *MYB162*; *PHOSPHOLIPID TRANSFER PROTEIN HOMOLOG1, PLT1*; and *PLT2*) were associated with WEs 47:0, 49:0, and/or 54:0.

### Hotspot analysis for waxes

Given that biosynthetic pathways and regulatory networks can be shared by wax compounds (Post-Beittenmiller, 1996; Lewandowska *et al*., 2020), we sought to identify genomic hotspots that possessed GWAS signals for several wax traits, potentially housing genes involved in the deposition of multiple cuticular waxes. Through the implementation of a sliding window method, we identified 34 and 15 GWAS hotspots for traits within (intraclass) and across (interclass) wax classes, respectively (Figure 3; Table S12). With the exception of FA, each of the wax classes had at least one intraclass hotspot. The PA and WE classes had the highest number with 10 and 20 intraclass hotspots, respectively. Notably, 65% of the WE hotspots were associated with one or more of the five WEs (47:0, 49:0, 50:0, 52:0, 54:0) found to be most important for predicting *g*_c_ in the RF analysis. Of the 15 interclass hotspots, nine were coincident (ie., at least 50% overlapping intervals) with 12 of the intraclass hotspots (Figure 3; Table S12). Among the 15 interclass hotspots identified, six were associated with one to five WEs, along with one to six additional traits drawn from up to two other wax classes (AD, PA, HC, and/or FA).

**Figure 3.**
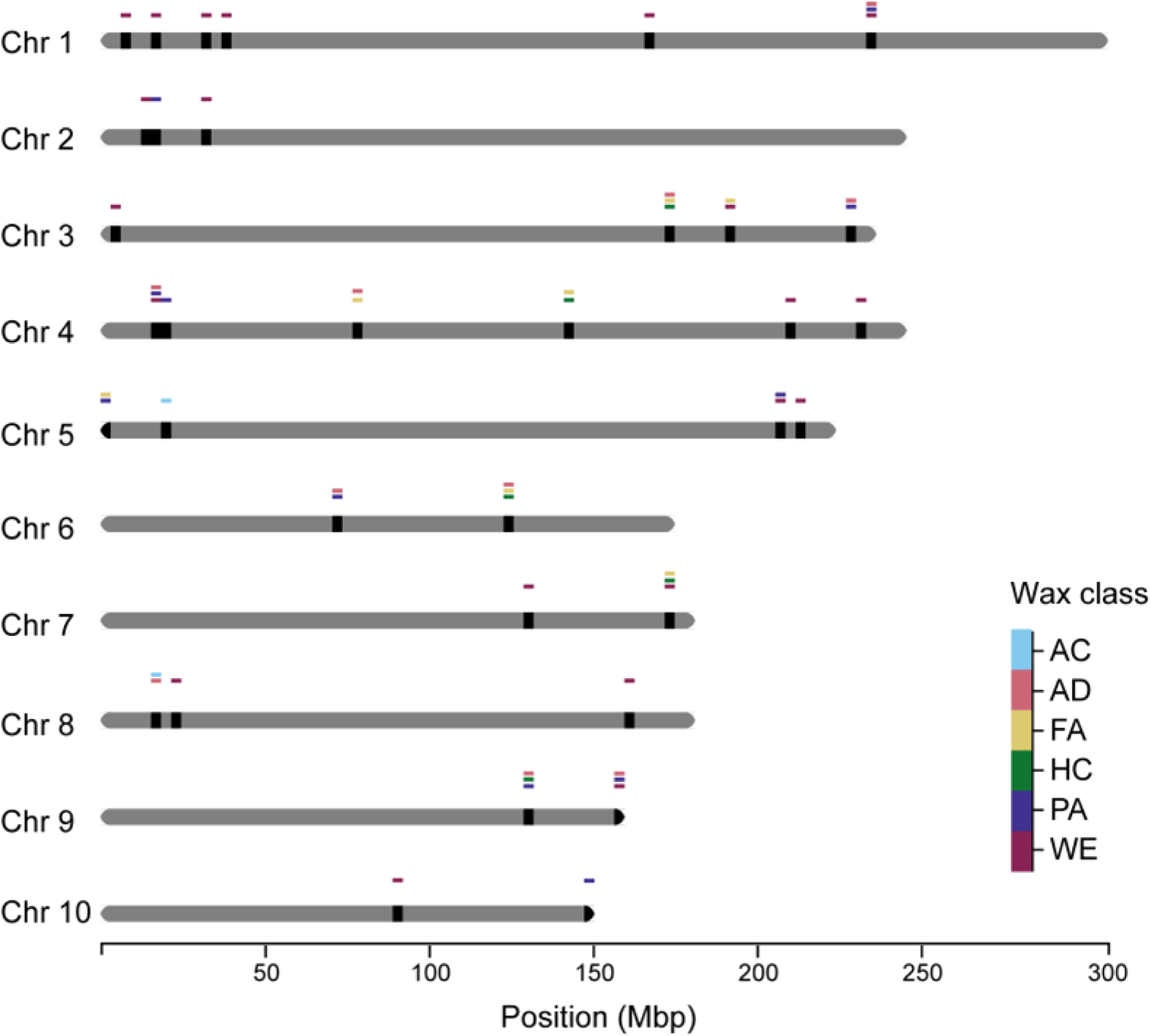
Genomic positions of GWAS hotspots in the maize genome. A chromosomal representation of GWAS hotspots was generated by plotting their physical positions (Mb, B73 RefGen_v4) across the 10 chromosomes of maize (x-axis). On each chromosome, black rectangles indicate the positions of GWAS hotspots. Hotspots are depicted at the wax class level and color-coded as follows: Alicyclic compounds (AC; pale blue), Aldehydes (AD; pink), Fatty acids (FA; yellow), Hydrocarbons (HC; green), Primary alcohols (PA; purple), and Wax esters (WE; bordeaux). Hotspots with stacked color-coded bars represent interclass hotspots, while those with a single color-coded bar represent intraclass hotspots.

The combined set of intra- and interclass GWAS hotspots contained 109 unique loci associated with 45 traits that spanned all six wax classes. Collectively these hotspots contain 371 unique genes. Of these, 33 were also identified by TWAS, and 110 by FCT, but not necessarily for the same traits associated via GWAS (Table S13). Twenty-one of the hotspot genes are among the 231 higher confidence candidates identified by all three methods (Tables S13 and S17), of which three are listed as plausible candidates in Table 1. The three plausible and higher confidence candidate genes located in GWAS hotspots are as follows: *Zm00001d013798* (CAS1 homolog) in an intraclass hotspot for ACs; *PAT38* in an intraclass hotspot for PAs; and *Zm00001d049100* (TOL homolog) in intraclass hotspots for PAs and WEs (Tables 1, S14, S16, and S17). Additionally, the TOL homolog *Zm00001d049100* was located in the most trait-dense interclass hotspot we identified, which was associated with four PA, one AD, and five WE traits, with four of these WEs (47:0, 49:0, 50:0, 52:0) identified through RF analysis as strongly predictive of *g*_c_. Notably, wax traits associated with each of these Group 1 genes by all three methods are among those associated with the intraclass hotspots in which the genes reside (ACs for the CAS1 homolog *Zm00001d013798*; PAs for PAT38 and the TOL homolog; see Table S10).

We identified an additional set of four plausible candidate genes located within hotspots according to the relevance of their predicted functions to the abundance of cuticular waxes (Table S13). Although these five genes were not found by TWAS, they were found associated with one or more wax traits by both GWAS and FCT. A second gene (*Zm00001d008671*) encoding a predicted CAS1 homolog was found within an intraclass hotspot for ACs and an interclass hotspot for AC and AD traits on chromosome 8 (Table S13). Interestingly, within an intraclass hotspot for PAs on chromosome 1, two genes (*Zm00001d032719* and *Zm00001d032721*), approximately 106 kb apart, encode homologs of FATTY ALCOHOL OXIDASE4b (FAO4b) found in Arabidopsis (Table S13). Lastly, *Zm00001d049155*, a second associated SEC23/SEC24 homolog, was located in an intraclass hotspot for PAs.

### Investigating a shared genetic basis between waxes and *g*_c_

Given that the RF analysis showed a moderate predictability of *g*_c_ with all 60 waxes, we integrated the GWAS (0.002% SNPs), TWAS (top 0.25% genes), and FCT (top 0.25% genes) results from both this study and Lin *et al*. (2022) to explore the extent to which natural variation in leaf cuticular waxes and *g*_c_ is connected through a shared underlying genetic architecture. A total of 17 unique candidate genes were identified in GWAS, 9 in TWAS, and 10 in FCT for both *g*_c_ and at least one wax trait (Table S14). Collectively, this set of 35 non-redundant genes was associated with 28 unique wax traits spanning all six wax classes. Among the 35 genes, only *Zm00001d017380*, encoding a homolog of Arabidopsis MEMBRANE STEROID BINDING PROTEIN1, was detected by more than one method (TWAS and FCT). The set of 35 genes included four identified by Lin *et al*. (2022) as plausible candidates associated with *g*_c_: *Zm00001d038404*/*Zm00001d038405* (encodes a Ypt/Rab-GAP homolog) for PA 36:0; *Zm00001d012175* (encodes PECTIN ACETYLESTERASE 5, PAE5) for HC 27:0, *Zm00001d005087* (encodes G2-LIKE-TRANSCRIPTION FACTOR 35, GLK35) for AD 30:0, total AD, AC β-Amyrin, and total wax; and *Zm00001d050185* (encodes MYB108) for HC 35:0. Collectively, the predicted functions of proteins encoded by these genes implicate the importance of cell wall biosynthesis (PAE5), transcriptional regulators of cuticle development (GLK35 and MYB108), and intracellular membrane trafficking (Ypt/Rab-GAP homolog) in the genetic control of both *g*_c_ and waxes.

## DISCUSSION

Waxes are critical for the cuticle function in protecting plants from biotic and abiotic stresses, including drought stress. Most prior studies identifying genetic determinants of cuticle biogenesis in maize and other grasses have focused on juvenile (seedling) leaves. However, the life cycle of maize is dominated by adult leaves, whose cuticular wax profiles are very different from juvenile leaves (Bourgault *et al*., 2020). In this study, we leveraged natural variation in the abundance of 60 wax traits scored on adult maize leaves belonging to 310 diverse inbred lines of the WiDiv panel. Our overarching goal was to search for genetic determinants of adult cuticular wax accumulation and to analyze the relationship between wax composition and *g_c_* to determine which waxes are most important for protecting adult leaves from water loss across the cuticle.

The prevailing consensus is that the water barrier property of the cuticle is conferred mainly by waxes (Schönherr, 1976; Kerstiens, 2006; Isaacson *et al*., 2009; Jetter and Riederer, 2016) and that wax composition, rather than quantity, is critical to this function (Vogg *et al*., 2004; Jetter *et al*., 2018; Buschhaus and Jetter, 2012; Jetter and Riederer, 2016). Consistent with that view, the RF analysis demonstrated an overall predictability of 0.27 for *g*_c_ based on all 60 cuticular waxes combined (Figure 1; Table S3). Significant correlations were observed between *g*_c_ and variation in the abundance of 18 individual waxes (Figure S1; Table S2), including six compounds identified as most critical for predicting *g*_c_ in the RF analysis (Figure 1; Table S3): five high molecular weight WEs (47:0, 49:0, 50:0, 52:0, 54:0) and FA 22:0. This extends similar findings from an earlier RF analysis employing a subset of the inbred lines used here (Lin *et al*., 2022) and is consistent with findings implicating WEs with carbon chains > 48:0 in the establishment of water barrier properties of the cuticle during adult maize leaf development (Bourgault *et al*., 2020). The longer the carbon chain, the higher their melting point, and consequently, their hydrophobicity (Patel *et al*., 2001), potentially explaining the importance of these WEs for waterproofing of the cuticle. Consistent with our findings, increased water permeability has been shown for Arabidopsis *wax ester synthase1* (*wsd1*) mutants with reduced WE load (Patwari *et al*., 2019). Yet, our finding that the majority of *g*_c_ variation is not predicted by total wax content demonstrates that other factors, such as the relative abundances of wax components, are also important for the water barrier function of the cuticle.

To elucidate the genetic architecture of cuticular wax accumulation, the abundances of 60 wax traits were analyzed as quantitative traits in GWAS employing ∼10 million SNPs, in TWAS investigating associations with transcript abundances of ∼20,000 genes expressed in cuticle maturation zones, and in FCT combining results of GWAS and TWAS. Intersectional approaches were employed to identify genes whose potential roles in cuticular wax accumulation are supported by multiple lines of evidence. To enhance this effort, we also identified genomic hotspots as loci associated via GWAS with more than one wax trait, and compared lists of candidate genes identified in this study with those identified in our earlier studies searching for genetic determinants of *g*_c_ and cuticle development.

Among the 231 higher confidence candidate genes identified by GWAS, TWAS, and FCT (Table S9), six are featured in Table 1 as plausible candidate genes based on predicted functions in cuticular wax biosynthesis. One of these, *CER8* (*Zm00001d024723*), encodes a LACS with a previously demonstrated role in the accumulation of epicuticular waxes of juvenile leaves through mutant analysis (Zheng *et al*., 2019). LACS enzymes convert C16 and C18 fatty acids into activated CoA thioesters, which are precursors of all aliphatic waxes (Yeats and Rose, 2013). Consistent with a broad role for LACS enzymes in wax biosynthesis, our study detected an association of maize *CER8* with members of both the acyl reduction (PA 32:0) and alkane-forming pathways (HC 37:0) (Table S9).

KCS enzymes are components of FAE complexes that function downstream of LACS enzymes to elongate C16 and C18 CoA thioesters, forming VLCFAs and their derivatives, which comprise the majority of cuticular wax (Lee and Suh, 2013). We identified two KCS enzyme genes as higher confidence candidates associated with a variety of wax traits by all three methods (Table 1): *KCS22* (*Zm00001d045660*) and *GL4B/KCS5* (*Zm00001d017941*). While neither gene has an experimentally validated function in wax biosynthesis, *GL4A,* a paralog of *GL4B*, has a demonstrated role in the accumulation of juvenile epicuticular waxes (Liu *et al*., 2009). *GL4B* was associated with five WEs and FA 24:0. *KCS22* was associated with more wax traits than any other gene in our study (26 altogether) and is one of the most highly associated genes for many of these traits (Table S9). The associations we observed for both *KCS* genes with waxes that are products of the acyl reduction and alkane-forming pathways are consistent with the established functions of KCS enzymes.

Two genes encoding GDSL lipases were identified as higher confidence candidates (Table 1), together associating with six wax traits belonging to three classes (FA, HC and WE; Table S9). *GELP2* (*Zm00001d018474*), with expression in stomata (Sun *et al*., 2022), encodes a homolog of Arabidopsis *CUS2*, a putative cutin synthase promoting the accumulation of cutin polymers and cuticular ridges in sepals (Hong *et al*., 2017). *GELP2* may therefore impact wax accumulation indirectly via a function in cutin matrix assembly. The other GDSL lipase gene we identified, *Zm00001d046642,* is a homolog of *Arabidopsis OSP1*, which is required for stomatal cuticular ledge formation but does not affect cutin accumulation (Tang *et al*., 2020). Biochemically, OSP1 functions as a thioesterase catalyzing the formation of VLCFAs and promoting the accumulation of all classes of aliphatic cuticular waxes (Tang *et al*., 2020). Thus, the maize *OSP1* homolog we identified could impact wax biosynthesis directly as well.

Another class of wax biosynthetic enzyme genes identified in our study is represented by a closely spaced pair of *FAO4b* genes (*Zm00001d032719* and *Zm00001d032721*) . These were not associated with wax traits by all three methods, but are both located in an intraclass GWAS hotspot for PAs (Table S13). Both genes were associated with PAs via FCT as well, and are thus well-supported as candidate determinants of PA biosynthesis. Recent work on FAOs in *Arabidopsis* demonstrated a function for two FAO enzymes in the oxidation of PAs, converting them to ADs (Yang *et al*., 2022). Thus, our finding of an association of two *FAO4b* genes with variation in PA accumulation is consistent with what is known about the functions of FAOs in *Arabidopsis*.

AC waxes (triterpenes and steroids) are synthesized independently from VLCFA-derived aliphatic waxes, via the mevalonate pathway involving oxidosqualene cyclases (OSCs) that convert 2,3-oxidosqualene into steroids and triterpenes (Thimmappa *et al*., 2014). A putative OSC gene identified by all three methods in our study and listed in Table 1 is *Zm00001d013798*, encoding a homolog of the *Arabidopsis* OSC gene *CAS1*. *Zm00001d013798* was associated with multiple AC waxes via all three methods and is the first or second most highly associated gene for the majority of these wax traits via both TWAS and FCT (Table S9). This gene is also located in an intraclass GWAS hotspot on chromosome 5 for multiple AC waxes (Table S13). Another *Arabidopsis* CAS1 homolog, *Zm00001d008671*, was identified in a different GWAS hotspot on chromosome 8 for AC waxes (total AC, α-amyrin, and β-amyrin). While not associated with ACs or other wax traits via TWAS, *Zm00001d008671* was also highly associated with total ACs, α-amyrin, and β-amyrin via FCT (Table S13). Both OSC genes we identified are therefore strongly supported as key genes for AC wax biosynthesis.

Based on the established role of membrane trafficking in the transport of waxes to the cell surface (McFarlane *et al*., 2014), we have highlighted a variety of potential intracellular trafficking proteins among our candidate genes for maize cuticular wax biogenesis. SEC23/SEC24 proteins initiate COPII-mediated vesicle transport between the ER and the Golgi apparatus (Brandizzi and Barlowe, 2013; Cui *et al*., 2022). *Zm00001d013301*, which encodes a SEC23/SEC24 homolog, is a higher confidence candidate gene associated with one FA via TWAS and FCT, and with HCs via GWAS and FCT (Tables 1 and S13). This gene was named *SAG4* (*SALT-TOLERANCE-ASSOCIATED-GENE4*) and confers salt tolerance in maize (X., Luo *et al*., 2019), but has not been previously linked to cuticles. Another SEC23/24 gene, *Zm00001d049155*, is located in a GWAS hotspot for PAs and was also associated with PA via FCT, although not via TWAS ( Table S13). These associations suggesting a role for COPII-mediated vesicle trafficking in wax accumulation are of particular interest in relation to a recent finding that a mutation in a *SEC23* gene in cucumber causes a glossy phenotype and drastically reduces cuticular wax load in fruits (Gao *et al*., 2023).

The exocyst complex functions at the plasma membrane as vesicle tether, facilitating vesicle fusion at appropriate sites on the cell surface (Cui *et al*., 2022). We found a gene encoding a putative exocyst subunit EXO84B (*Zm00001d019104*) to be associated with FA 32:0 by GWAS, and with HC 39:0 by TWAS and FCT (Tables 1 and S13), suggesting a role for the exocyst complex in vesicle-mediated delivery of these waxes to the outer cell surface. Another notable candidate gene with a predicted function in vesicle targeting is (*Zm00001d038404/Zm00001d038405*), encoding a Ypt/Rab-GAP domain family protein (Table 1). This gene was associated with several high molecular weight WEs by TWAS and FCT; it was the most strongly associated gene by TWAS for WEs 54:0 and 56:0 (Table S9). This gene is of particular interest because it was identified as a plausible candidate determinant of *g*_c_ in our prior GWAS (Lin *et al*., 2022), and because four of the WEs associated with this gene via TWAS and/or FCT (49:0, 50:0, 52:0, and 54:0) are among the top six compounds most predictive of *g*_c_ (Figure 1). Taken together, our findings suggest that this putative Ypt/Rab-GAP impacts *g*_c_ by playing a role in the transport of high molecular weight WEs. Rab GTPases located on vesicle surfaces direct vesicle trafficking through interactions with a variety of molecules including tethering proteins on target membranes; GAPs modulate their activity (Cui *et al*., 2022). Hence, a Ypt/Rab-GAP protein is predicted to play a role in vesicle targeting, possibly regulating a Rab GTPase that interacts with the exocyst complex as a membrane tether (Stenmark, 2009; Fendrych *et al*., 2013).

S-acylation is a post-translational modification mediating protein association with membranes (Batistič *et al*., 2008), potentially playing a role in localization of many proteins participating in cuticle delivery to and transport across the plasma membrane. We identified an S-acyltransferase gene, *Zm00001d026600 (PAT38;* Yuan *et al*., 2013), that is located in an intraclass GWAS hotspot for PAs (Table S13), and is also associated with PAs by TWAS and FCT; it was among the most highly associated genes by these two methods for total PA and several individual PAs (Tables 1 and S13). Thus, *PAT38* is strongly supported as a key determinant of PA accumulation through an unknown mechanism involving protein S-acylation. Interestingly, Yuan *et al*. (2013) showed that the expression of *PAT38* is responsive to osmotic stress, suggesting a possible role for *PAT38* in mediating environmental signals and modulating cuticular wax accumulation.

Another higher confidence candidate gene with strong, intersectional support, *Zm00001d049100*, encodes a TOL family protein with a potential function in intracellular trafficking of cuticle lipids. *Zm00001d049100* is located in a trait-dense interclass GWAS hotspot for multiple PAs and WEs, and one AD (Table S13); it was also strongly associated with multiple waxes belonging to the same three classes by TWAS and FCT. TOL family proteins in Arabidopsis are recognized for their ubiquitin-binding role initiating the signal for cargo protein entry into the multivesicular body (MVB) pathway (Winter and Hauser, 2006). While TOL proteins have not been previously linked to cuticle formation, the MVB pathway has been implicated in plant cuticle formation in Arabidopsis and maize (Goodman *et al*., 2021; Lin *et al*., 2022). Our findings for *Zm00001d049100* suggest a role for this gene in membrane trafficking processes important for accumulation of multiple aliphatic wax classes.

A final candidate gene of particular interest, *GLK35* (*Zm00001d005087*), encodes a MYB domain-containing putative transcription factor. It was identified in this study only by TWAS, but is notable because it was previously identified as a plausible candidate determinant of *g*_c_ via both TWAS and FCT (Lin *et al*. 2022; Table S18). Remarkably, *GLK35* was the gene most strongly associated with *g*_c_ via TWAS, underscoring its probable influence on *g*_c_, albeit through an impact on other waxes besides WEs, as indicated by the associations found in the present study via TWAS (with AC β-Amyrin, AD 30:0, total AD, and total wax). Notably, the two closest relatives of *GLK35* in *Arabidopsis* are genes encoding the recently characterized MYB-SHAQKYF transcription factors, MYS1 and MYS2 (Lin *et al*., 2022; Liu *et al*., 2022). These redundant transcriptional repressors directly suppress the expression of another transcriptional regulator of wax biosynthesis, *DECREASE WAX BIOSYNTHESIS*. Analysis of *mys1mys2* double mutants demonstrates that these genes promote wax biosynthesis (especially of HCs, the most abundant wax class in *Arabidopsis* leaves), protect leaves from water loss in a detached leaf assay, and confer drought tolerance (Liu *et al*., 2022). The relationship to *MYS1* and *MYS2* strengthens the hypothesis that *GLK35* acts as a determinant of *g*_c_ via regulation of wax biosynthesis genes.

With the exception of *CER8* (*Zm00001d024723*; Tables 1 and S13), encoding a putative LACS enzyme discussed earlier, none of our 231 higher confidence candidate genes (Table S9) have been previously linked by genetic evidence to cuticles. Thus, none of our higher confidence candidates correspond to *GLOSSY* genes with known identities. However the *GL26* gene (*Zm00001d008622*), encoding a FAE complex component (ECR/enoyl-CoA reductase; Fan, 2007), was associated with WE 43:0 via GWAS (Table S11) and with AD 28:0 via FCT (rank 45; Table S7). It is also among the 39 candidates listed in Table S11 based on overlap between genes associated with at least one trait in our study vs. top candidates for regulation of cuticle development identified by Qiao *et al*. (2020) via transcriptome analysis. The two *KCS* genes we identified as higher confidence candidates (*KCS5*/*GL4B* and *KCS22*; Table 1) are different from the three *KCSs* with genetic evidence for a function in cuticular wax accumulation (*KCS23/GL4*, Liu *et al*. 2009; *KCS19/AD1*, Liu *et al*. 2021; and *KCS12*, Xu *et al*., 2024), and also different from another *KCS* gene (*KCS24*) found to be associated with cuticle-dependent traits via GWAS by Xu *et al*. (2024). This lack of correspondence most likely reflects our focus on adult leaves, whereas all these other studies focus on juvenile leaves, whose cuticular wax profiles are quite different (Bourgault *et al*., 2020). Given the importance of adult leaves for agronomically significant traits, we believe that our study has identified many new genes that can ultimately be of value for breeding or engineering maize plants with improvements in cuticle-dependent traits of agronomic value.

## EXPERIMENTAL PROCEDURES

### Plant materials and experimental design

Cuticular wax data were collected from a set of 323 maize inbred lines from the Wisconsin Diversity (WiDiv) panel (Hansey *et al*., 2011) planted in two field environments at the University of California San Diego, San Diego, CA, in 2018. Line selection and field layout have been described in Lin *et al*. (2022).

### Evaluation and analysis of cuticular waxes and *g*_c_

Cuticular waxes were evaluated for the WiDiv panel using fully expanded leaves about one week after pollen shed. The middle intact section (15 cm; with no obvious pathogen or other damage) was sampled from the primary ear leaf (or one leaf immediately above/below) of four plants from each plot. Collected leaf tissues were wrapped in wet paper towels and shipped to Algoma University, Canada, on wet ice for wax analysis. Upon arrival, samples were stored at -20 °C until use. For analysis, leaves were thawed overnight at 4 °C, excess water removed by gently pressing both sides of the leaf on a paper towel, and dissected with a scalpel to obtain an 8-cm-long segment from the middle portion, removing the midrib. The resulting two pieces were then photographed alongside a ruler to determine surface area using ImageJ software (Abràmoff *et al*., 2004). Cuticular waxes were extracted by submerging the mature leaf tissue in chloroform for 60 s with gentle agitation. Three internal standards —n-tetracosane (24:0 alkane), 1-pentadecanol (15:0-OH) and heptadecanoic acid (17:0)— were added to the extract (5 μg each). Extracted wax samples were evaporated under a gentle stream of nitrogen, derivatized to form trimethylsilyl (TMS) ester and ether derivatives, resuspended in hexanes and analyzed by gas chromatography with flame ionization detection (GC-FIDs; two Thermo Scientific TRACE 1300 GC-FID systems were used) as described in Bourgault *et al*. (2020).

Cuticular waxes with missing values in over 40% of the population, primarily due to levels falling below the minimum detection limits of GC-FID, were excluded from the analysis to mitigate the risk of biased results (Lubin *et al*., 2004). For each of the retained wax traits, abundance (µg·dm^-2^) was approximated for missing values by imputing a uniform random variable ranging from 0 to the minimum GC-FID detection value specific to the given trait within each environment (Lubin *et al*., 2004; Lipka *et al*., 2013).

To screen cuticular wax traits for significant outliers, a mixed linear model was fitted for each trait in ASReml-R version 3.0 (Gilmour *et al*., 2009). The model included the grand mean and check as fixed effects and genotype, environment, genotype-by-environment (G×E) interaction, incomplete block nested within environment, plot grid column nested within environment, GC-FID instrument, and GC-FID column nested within GC-FID instrument as random effects. The Studentized deleted residuals (Neter *et al*., 1996) generated from the mixed linear model were used to identify and remove outliers at a Bonferroni corrected threshold of α = 0.05. For each outlier-screened wax trait, the above model was fitted to estimate the variance components for calculating heritability on a line-mean basis (Table S1) following Lin *et al*. (2020). A best-fit model (Table S15) was selected for each outlier-screened phenotype based on an iterative mixed model fitting procedure in ASReml-R version 3.0 from which best linear unbiased predictors (BLUPs) were generated for each wax trait (Table S16). The BLUPs for *g*_c_ and flowering time (days to anthesis, DTA; Table S16) were obtained from a previous study of the identical subset of the WiDiv panel in the same two San Diego environments (Lin *et al*., 2022). The degree of association between BLUPs of wax traits and *g*_c_ was estimated using the Pearson’s correlation coefficient with the ‘cor.test’ function in R version 3.5.1 (R Core Team, 2019).

### Associations between cuticular wax traits and *g*_c_

To assess the predictive ability of *g*_c_ using 60 cuticular wax traits, random forest (RF) models were fitted using 310 inbred lines in the WiDiv panel following that of Lin *et al*. (2022). Briefly, forests were grown with the ‘cforest’ function from the R package ‘party’ version 1.3-7 (Strobl *et al*., 2007). Fivefold cross-validation was performed 50 times to evaluate the mean predictive accuracy for *g*_c_ as described in Lin *et al*. (2020). Variable importance measures were calculated using the ‘varimp’ function in the ‘party’ package. Parameters used for the number of trees grown (*ntree* = 1000) and predictors sampled (*mtry* = 10) were those that maximized predictive accuracy.

### Genomic data sets for association analyses

A set of 9,715,072 SNPs in B73 RefGen_v4 coordinates (Lin *et al*., 2022) for 310 lines with cuticular wax values were used to conduct GWAS. Briefly, a reference SNP genotype set (Wu *et al*., 2021) derived from maize HapMap 3.2.1 (Bukowski *et al*., 2018) was imputed based on a target GBS SNP set via BEAGLE v5 (Browning *et al*., 2018). Biallelic SNPs with MAF ≥ 5% and predicted dosage *r*^2^ (DR2) ≥ 0.80 were retained for GWAS (Data S1).

Transcript abundance for 20,013 genes across 310 lines (Lin *et al*., 2022) were used for TWAS. The Lexogen QuantSeq 3′ mRNA-sequencing data were generated from the proximal, immature and actively growing section of unexpanded adult leaves (sheath length < 2 cm). For each gene, BLUPs were calculated to combine expression levels from two environments in a mixed linear model. The probabilistic estimation of expression residuals (PEER) (Stegle *et al*., 2010) approach was applied to the matrix of BLUP expression values to account for inferred confounders. The resultant PEER values after extracting 20 latent factors were further filtered for outliers using Studentized deleted residuals and then used for conducting TWAS (Data S2).

### Genome-wide association study

To reduce heteroscedasticity and nonnormality of residuals in association tests, the Box-Cox power transformation (Box and Cox, 1964) was applied to each wax trait with an intercept-only model to choose the optimal value of convenient lambda (Table S17) for transforming the BLUP values with the MASS package version 7.3-51.4 in R version 3.5.1 (R Core Team, 2019). An additional outlier removal was performed using Studentized deleted residuals to generate the outlier-screened transformed BLUP values (Figure S2; Table S18) of wax traits for association analyses.

With the outlier-screened transformed BLUP values of each cuticular wax trait, associations were tested with each of the 9,715,072 SNPs using a mixed linear model (Zhang *et al*., 2010) in the R package GAPIT version 3.0 (Lipka *et al*., 2012) according to Lin *et al*. (2022). The mixed linear model included days to anthesis BLUPs, principal components based on SNP genotype data, and a kinship matrix to control for variation in maturity, population stratification, and unequal relatedness, respectively. Principal components and the kinship matrix were calculated as described in Lin *et al*. (2022). The optimal model for GWAS selected based on the Bayesian information criterion (Schwarz, 1978) included only the kinship matrix for nearly all wax traits, except for the models of two traits (fatty acid C:20 and fatty acid C:22) that also included days to anthesis as a covariate. A likelihood-ratio-based *R^2^* statistic (*R^2^* _LR_) (Sun *et al*., 2010) was used to approximate the amount of phenotypic variation explained by a SNP for a transformed wax trait.

### Transcriptome-wide association study

For each of the wax traits (outlier-screened transformed BLUPs), TWAS was performed with PEER values for each of the 20,013 expressed genes to identify strong associations between trait and gene expression levels. A mixed linear model with the same kinship matrix from GWAS was fitted using the *gwas* function with the P3D function set to FALSE in the R package ‘rrBLUP’ version 4.6 (Endelman, 2011).

### Fisher’s combined test

To integrate GWAS and TWAS results, the Fisher’s combined test (FCT) was performed as described in Kremling *et al*. (2019). Briefly, for each wax trait, the top 10% most associated SNPs (971,508) from GWAS were assigned to the nearest gene. The *P*-values of genes not tested in TWAS were set to 1. The paired GWAS and TWAS *P*-values for each gene were used to conduct a FCT with the sumlog method in the R package ‘metap’ version 1.4 (Dewey, 2016).

### Candidate gene identification

Because of the different statistical power and model structure in GWAS, TWAS and FCT, potential candidate genes were selected based on the rankings of *P*-values for each statistical method as described in Kremling *et al*. (2019). To identify candidate genes from the GWAS results, the top 0.002% of SNPs were used to declare loci associated with each wax trait following that of Wu *et al*. (2021). Given that genome-wide LD decayed to nominal levels by ∼200 kb in the WiDiv panel (Lin *et al*., 2020), a ± 200 kb search interval centered by the peak SNP for each declared locus was applied for candidate gene identification according to Lin *et al*. (2020). For each wax trait, the top 0.25% associated genes were selected from TWAS and FCT results based on their *P*-values. On average, GWAS, TWAS and FCT identified comparable numbers of candidate genes for each wax trait.

### Identification of GWAS hotspots

To identify genomic regions containing multiple GWAS signals for several wax traits or “GWAS hotspots,” genome-wide scans were performed with the set of GWAS-identified loci within (5-15 traits) and across (59 traits not including total wax) six wax trait classes: primary alcohol (PA; 7 compounds and 1 sum trait), fatty acid (FA; 11 compounds and 1 sum trait), hydrocarbon (HC; 9 compounds and 1 sum trait), aldehyde (AD; 4 compounds and 1 sum trait), wax ester (WE; 14 compounds and 1 sum trait), and alicyclic (AC; 8 compounds and 1 sum trait). To generate a reference distribution for the number of detected GWAS loci across the genome, we implemented a sliding window procedure that used a window size of 200 kb and step size of one-fifth of the window size to approximate the number of loci (i.e., peak SNPs) within (intraclass) and across (interclass) the six wax trait classes. A 95% quantile of the resultant distributions indicated that an intraclass GWAS hotspot should contain at least three GWAS-identified loci, and an interclass GWAS hotspot should have at least four GWAS-identified loci. The physical boundaries of a GWAS hotspot were defined based on the entirety of all partially overlapping windows that satisfied the above criteria. A hotspot was considered to be interclass only if it included traits from two or more wax classes. Window sizes of 10 and 50 kb produced nearly equivalent results. The positions of GWAS hotspots in the maize genome were plotted in the R package ‘ChromoMap’ version 4.1.1 (Anand and Rodriguez Lopez, 2022).

## AUTHOR CONTRIBUTIONS

M.A.G., I.M., L.G.S., and M.J.S. conceived the research plans and supervised the experiments. S.M. and M.V. managed the fields, collected tissue samples, and generated and analyzed data for cuticular conductance. R.B., M.M., and S.V.B. performed the analysis of cuticular wax. P.Q. conducted the analysis of 3′ mRNA-seq data. M.L. and H.B. conducted the quantitative genetic analyses and wrote the first draft of the article, with input from all authors. M.A.G. agrees to serve as the author responsible for contact and ensures communication.

## ACKNOWLEDGEMENTS

We especially thank Akriti Bhattarai, a BTI intern student, Albert Nguyen, Lesley Saldana De Haro, Alfredo Arriola, Hiep Ha, Jessica Davis, Cameron Garland, Anasilvia Herrera Fuentes, Maria Fernanda Salcedo, and Alondra Deras at UCSD for collecting phenotypic data and tissue samples. We also thank Elise Withers, a BTI intern student, for preliminary evaluation of an early generation computational pipeline for conducting TWAS, and to Amanda Dubois, Annika Sonntag, Aseel Hashim, Caleb Charlebois, Elvis Boakye, and Nora Sonntag for help with extracting wax samples and calculating leaf areas.

## DATA AVAILABILITY STATEMENT

All raw 3′-mRNA-seq data are available from the NCBI Sequence Read Archive under BioProject PRJNA773975. Supporting Data S1–2 are available on CyVerse (https://datacommons.cyverse.org/browse/iplant/home/shared/GoreLab/dataFromPubs/Lin_ <u>CuticularWaxTWAS_2024</u>). Code is available from Github (https://github.com/GoreLab/Maize_Cuticular_Wax).

## SUPPORTING INFORMATION

**Figure S1.** Correlation matrix for best linear unbiased predictors (BLUPs) of maize adult leaf cuticular conductance (*g*_c_; g·h^-1^·g^-1^) and wax composition (µg·dm^-2^). Pearson’s correlation coefficients (r) are presented in the upper right triangle, while the corresponding P-values for the significance of associations (α = 0.05) are displayed below the diagonal. Waxes of the same class are highlighted by red rectangles and brackets.

**Figure S2.** Box plots of transformed best linear unbiased predictors (BLUPs) of cuticular waxes (µg·dm^-2^) in the maize Wisconsin diversity panel. (A) Box plots showing the abundance of each wax compound. (B) Box plots showing the total abundance of each wax class, including PA (primary alcohol), FA (fatty acid), HC (hydrocarbon), AD (aldehyde), WE (wax ester), and AC (alicyclic compound). Box limits indicate the upper and lower quartiles; center lines in boxes indicate the median value; whiskers indicate 1.5x interquartile range; and dots indicate outliers.

**Table S1.** Means, ranges, and variances for best linear unbiased predictors (BLUPs) of adult maize leaf cuticular waxes (PA: primary alcohol; FA: fatty acid; HC: hydrocarbon; AD: aldehyde; WE: wax ester; AC: alicyclic; µg·dm^-2^) for 310 maize inbred lines from two environments in San Diego in 2018, and estimated heritabilities on a line-mean basis.

**Table S2.** Pearson’s correlation of *g*_c_ to cuticular wax composition. The upper triangle of the matrix shows Pearson’s correlations coefficients between best linear unbiased predictors for *g*_c_ and adult maize leaf cuticular waxes (PA: primary alcohol; FA: fatty acid; HC: hydrocarbon; AD: aldehyde; WE: wax ester; AC: alicyclic), whereas the lower triangle shows the significance of the correlations as P-values.

**Table S3.** Importance scores and ranks of adult maize leaf cuticular waxes (PA: primary alcohol; FA: fatty acid; HC: hydrocarbon; AD: aldehyde; WE: wax ester; AC: alicyclic) from random forest regression to predict cuticular conductance (*g*_c_) in 310 inbred lines of the Wisconsin diversity panel.

**Table S4.** Genomic information (B73 RefGen_v4) and association statistics for the top 0.002% SNPs associated with adult maize leaf cuticular waxes in a genome-wide association study in the Wisconsin diversity panel.

**Table S5.** Genomic information (RefGen_v4) for the candidate genes residing ± 200 kb of the peak SNPs associated with adult maize leaf cuticular waxes in a genome-wide association study in the Wisconsin diversity panel.

**Table S6.** Genomic information (RefGen_v4) for the top 0.25% associated genes for adult maize leaf cuticular waxes in a transcriptome-wide association study in the Wisconsin diversity panel.

**Table S7.** Genomic information (RefGen_v4) for the top 0.25% genes associated with adult maize leaf cuticular waxes in Fisher’s combined test in the Wisconsin diversity panel.

**Table S8.** Number of candidate genes identified from a genome-wide association study (GWAS), transcriptome-wide association analysis (TWAS), and Fisher’s combined test (FCT) of adult maize leaf cuticular waxes (PA: primary alcohol; FA: fatty acid; HC: hydrocarbon; AD: aldehyde; WE: wax ester; AC: alicyclic).

**Table S9.** Higher confidence candidate genes associated with adult maize leaf cuticular waxes in all three methods (GWAS, TWAS and FCT) and their grouping based on whether they were identified for the same trait or wax class.

**Table S10.** Rice and Arabidopsis homologs of plausible candidate genes identified for adult maize leaf cuticular waxes via a genome-wide association study (GWAS), transcriptome-wide association study (TWAS), Fisher’s combined test (FCT) and GWAS hotspot analysis in the maize Wisconsin diversity panel.

**Table S11.** Overlapping candidate genes for maize leaf cuticle biosynthesis and development regulation (Qiao et al. 2020) and maize adult leaf cuticular waxes.

**Table S12.** Genomic hotspots and loci (peak SNPs) associated with adult maize leaf cuticular waxes in a genome-wide association study in the Wisconsin diversity panel.

**Table S13.** Candidate genes associated with adult maize leaf cuticular waxes by GWAS hotspot analysis and their associated traits in TWAS and FCT.

**Table S14.** Overlapping candidate genes for maize adult leaf cuticular conductance (*g*_c_; Lin et al. 2022) and cuticular waxes.

**Table S15.** Best fitted models used to calculate best linear unbiased predictors (BLUPs) for adult maize leaf cuticular waxes (PA: primary alcohol; FA: fatty acid; HC: hydrocarbon; AD: aldehyde; WE: wax ester; AC: alicyclic) according to a likelihood ratio test (α = 0.05). The star (*) indicates that a random effect term was retained in the mixed linear model, whereas the ‘x’ indicates that a fitted random effect term was not significant and removed from the mixed linear model.

**Table S16.** Best linear unbiased predictors (BLUPs) of adult maize leaf cuticular conductance (*g*_c_; g·h^-1^·g^-1^), BLUPs of flowering time (DTA; days to anthesis) and BLUPs of cuticular waxes (PA: primary alcohol; FA: fatty acid; HC: hydrocarbon; AD: aldehyde; WE: wax ester; AC: alicyclic; µg·dm^-2^) for 310 maize inbred lines from two environments in San Diego in 2018.

**Table S17.** Lambda values used to transform best linear unbiased predictors (BLUPs) of adult maize leaf cuticular waxes (PA: primary alcohol; FA: fatty acid; HC: hydrocarbon; AD: aldehyde; WE: wax ester; AC: alicyclic; µg·dm^-2^) for 310 maize inbred lines from two environments in San Diego in 2018.

**Table S18.** Transformed best linear unbiased predictors (BLUPs) of adult maize leaf cuticular waxes (PA: primary alcohol; FA: fatty acid; HC: hydrocarbon; AD: aldehyde; WE: wax ester; AC: alicyclic; µg·dm^-2^) for 310 maize inbred lines from two environments in San Diego in 2018. Outlier values for each wax trait were replaced by NA.

**Data S1**. SNP marker genotypes used for GWAS and the kinship matrix used for both GWAS and TWAS.

**Data S2**. Transcript abundances used for TWAS.

## REFERENCES

Abràmoff, M., Magalhães, P.J. and Ram, S. (2004) Image processing with ImageJ. Biophotonics international, 11, 36–42.

Albersheim, P., Darvill, A., Roberts, K., Sederoff, R. and Staehelin, A. (2010) Plant Cell Walls, Garland Science.

Anand, L. and Rodriguez Lopez, C.M. (2022) ChromoMap: an R package for interactive visualization of multi-omics data and annotation of chromosomes. BMC Bioinformatics, 23, 33.

Babiychuk, E., Bouvier-Navé, P., Compagnon, V., Suzuki, M., Muranaka, T., Van Montagu, M., Kushnir, S. and Schaller, H. (2008) Allelic mutant series reveal distinct functions for *Arabidopsis* cycloartenol synthase 1 in cell viability and plastid biogenesis. Proc. Natl. Acad. Sci. USA, 105, 3163–3168.

Bargel, H., Barthlott, W., Koch, K., Schreiber, L. and Neinhuis, C. (2004) Plant cuticles: multifunctional interfaces between plant and environment. In A. R. Hemsley and I. Poole, eds. The Evolution of Plant Physiology. Academic Press, pp. 171–III.

Batistič, O., Sorek, N., Schültke, S., Yalovsky, S. and Kudla, J. (2008) Dual fatty acyl modification determines the localization and plasma membrane targeting of CBL/CIPK Ca2+ signaling complexes in Arabidopsis. Plant Cell, 20, 1346–1362.

Bianchi, G., Avato, P. and Salamini, F. (1979) Glossy mutants of maize. Heredity, 42, 391–395.

Bi, H., Kovalchuk, N., Langridge, P., Tricker, P.J., Lopato, S. and Borisjuk, N. (2017) The impact of drought on wheat leaf cuticle properties. BMC Plant Biol., 17, 85.

Bourgault, R., Matschi, S., Vasquez, M., et al. (2020) Constructing functional cuticles: Analysis of relationships between cuticle lipid composition, ultrastructure and water barrier function in developing adult maize leaves. Ann. Bot., 125, 79–91.

Box, G.E.P. and Cox, D.R. (1964) An analysis of transformations. J. R. Stat. Soc., 26, 211–243.

Brandizzi, F. and Barlowe, C. (2013) Organization of the ER–Golgi interface for membrane traffic control. Nat. Rev. Mol. Cell Biol., 14, 382–392.

Browning, B.L., Zhou, Y. and Browning, S.R. (2018) A one-penny imputed genome from next-generation reference panels. Am. J. Hum. Genet., 103, 338–348.

Bukowski, R., Guo, X., Lu, Y., et al. (2018) Construction of the third-generation *Zea mays* haplotype map. GigaScience, 7, 1–12.

Buschhaus, C., Herz, H. and Jetter, R. (2007) Chemical composition of the epicuticular and intracuticular wax layers on adaxial sides of Rosa canina leaves. Ann. Bot., 100, 1557–1564.

Buschhaus, C. and Jetter, R. (2012) Composition and physiological function of the wax layers coating Arabidopsis leaves: β-amyrin negatively affects the intracuticular water barrier. Plant Physiol., 160, 1120–1129.

Corey, E.J., Matsuda, S.P. and Bartel, B. (1993) Isolation of an Arabidopsis thaliana gene encoding cycloartenol synthase by functional expression in a yeast mutant lacking lanosterol synthase by the use of a chromatographic screen. Proc. Natl. Acad. Sci. USA, 90, 11628–11632.

Cui, L., Li, H., Xi, Y., et al. (2022) Vesicle trafficking and vesicle fusion: mechanisms, biological functions, and their implications for potential disease therapy. Mol Biomed, 3, 29.

DeBono, A., Yeats, T.H., Rose, J.K.C., Bird, D., Jetter, R., Kunst, L. and Samuels, L. (2009) *Arabidopsis* LTPG Is a glycosylphosphatidylinositol-anchored lipid transfer protein required for export of lipids to the plant surface. Plant Cell, 21, 1230–1238.

Dewey, M. (2016) metap: Meta-analysis of significance values. R package version 0.7.

Dietrich, C.R., Perera, M.A.D.N., D Yandeau-Nelson, M., Meeley, R.B., Nikolau, B.J. and Schnable, P.S. (2005) Characterization of two GL8 paralogs reveals that the 3-ketoacyl reductase component of fatty acid elongase is essential for maize (Zea mays L.) development. Plant J., 42, 844–861.

Domínguez, E., Heredia-Guerrero, J.A. and Heredia, A. (2011) The biophysical design of plant cuticles: an overview. New Phytol., 189, 938–949.

Eigenbrode, S.D. (2002) Attachment to plant surface waxes by an insect predator. Integr. Comp. Biol., 42, 1091–1099.

Elango, D., Xue, W. and Chopra, S. (2020) Genome wide association mapping of epi-cuticular wax genes in Sorghum bicolor. Physiol. Mol. Biol. Plants, 26, 1727–1737.

Endelman, J.B. (2011) Ridge regression and other kernels for genomic selection with R package rrBLUP. Plant Genome, 4, 250–255.

Fan, L. (2007) Map based candidate gene cloning and functional analysis of genes involved in VLCFAs synthesis. M.S. Iowa State University.

Fendrych, M., Synek, L., Pecenková, T., Drdová, E.J., Sekeres, J., Rycke, R. de, Nowack, M.K. and Zársky, V. (2013) Visualization of the exocyst complex dynamics at the plasma membrane of Arabidopsis thaliana. Mol. Biol. Cell, 24, 510–520.

Fernández, V., Guzmán-Delgado, P., Graça, J., Santos, S. and Gil, L. (2016) Cuticle structure in relation to chemical composition: re-assessing the prevailing model. Front. Plant Sci., 7, 427.

Gao, L., Cao, J., Gong, S., Hao, N., Du, Y., Wang, C. and Wu, T. (2023) The COPII subunit CsSEC23 mediates fruit glossiness in cucumber. Plant J., 116, 524–540.

Gilmour, A.R., Gogel, B.J., Cullis, B.R. and Thompson, R. (2009) ASReml user guide release 3.0. VSN International Ltd, Hemel Hempstead, UK.

Goodman, K., Paez-Valencia, J., Pennington, J., Sonntag, A., Ding, X., Lee, H.N., Ahlquist, P.G., Molina, I. and Otegui, M.S. (2021) ESCRT components ISTL1 and LIP5 are required for tapetal function and pollen viability. Plant Cell, 33, 2850–2868.

Goodwin, S.M. and Jenks, M.A. (2005) Plant cuticle function as a barrier to water loss. In M.A. Jenks and P.M. Hasegawa, ed. Plant Abiotic Stress. Blackwell Publishing Ltd, pp. 14–36.

Hansey, C.N., Johnson, J.M., Sekhon, R.S., Kaeppler, S.M. and Leon, N. (2011) Genetic diversity of a maize association population with restricted phenology. Crop Sci., 51, 704–715.

Hong, L., Brown, J., Segerson, N.A., Rose, J.K.C. and Roeder, A.H.K. (2017) CUTIN SYNTHASE 2 Maintains Progressively Developing Cuticular Ridges in Arabidopsis Sepals. Mol. Plant, 10, 560–574.

Isaacson, T., Kosma, D.K., Matas, A.J., et al. (2009) Cutin deficiency in the tomato fruit cuticle consistently affects resistance to microbial infection and biomechanical properties, but not transpirational water loss. Plant J., 60, 363–377.

Jenks, M.A., Joly, R.J., Peters, P.J., Rich, P.J., Axtell, J.D. and Ashworth, E.N. (1994) Chemically Induced Cuticle Mutation Affecting Epidermal Conductance to Water Vapor and Disease Susceptibility in Sorghum bicolor (L.) Moench. Plant Physiol., 105, 1239–1245.

Jenks, M.A., Tuttle, H.A., Eigenbrode, S.D. and Feldmann, K.A. (1995) Leaf Epicuticular Waxes of the Eceriferum Mutants in Arabidopsis. Plant Physiol., 108, 369–377.

Jetter, R., Kunst, L. and Samuels, A.L. (2018) Composition of plant cuticular waxes. In M. Riederer and C. Müller, eds. Annual Plant Reviews. Chichester, UK: John Wiley & Sons, Ltd, pp. 145–181.

Jetter, R. and Riederer, M. (2016) Localization of the transpiration barrier in the epi- and intracuticular waxes of eight plant species: Water transport resistances are associated with fatty acyl rather than alicyclic components. Plant Physiol., 170, 921–934.

Jin, S., Zhang, S., Liu, Y., Jiang, Y., Wang, Y., Li, J. and Ni, Y. (2020) A combination of genome-wide association study and transcriptome analysis in leaf epidermis identifies candidate genes involved in cuticular wax biosynthesis in Brassica napus. BMC Plant Biol., 20, 458.

Kerstiens, G. (1996) Cuticular water permeability and its physiological significance. J. Exp. Bot., 47, 1813–1832.

Kerstiens, G. (2006) Water transport in plant cuticles: an update. J. Exp. Bot., 57, 2493–2499. Available at: [Accessed December 15, 2023].

Kim, H., Lee, S.B., Kim, H.J., Min, M.K., Hwang, I. and Suh, M.C. (2012) Characterization of glycosylphosphatidylinositol-anchored lipid transfer protein 2 (LTPG2) and overlapping function between LTPG/LTPG1 and LTPG2 in cuticular wax export or accumulation in Arabidopsis thaliana. Plant Cell Physiol., 53, 1391–1403.

Kolattukudy, P.E. (1970) Plant waxes. Lipids, 5, 259–275.

Krauss, P., Markstadter, C. and Riederer, M. (1997) Attenuation of UV radiation by plant cuticles from woody species. Plant, Cell Environ., 20, 1079–1085.

Kremling, K.A.G., Diepenbrock, C.H., Gore, M.A., Buckler, E.S. and Bandillo, N.B. (2019) Transcriptome-wide association supplements genome-wide association in *Zea mays*. G3, 9, 3023–3033.

Lee, S.B., Go, Y.S., Bae, H.-J., et al. (2009) Disruption of glycosylphosphatidylinositol-anchored lipid transfer protein gene altered cuticular lipid composition, increased plastoglobules, and enhanced susceptibility to infection by the fungal pathogen *Alternaria brassicicola*. Plant Physiol., 150, 42–54.

Lee, S.B. and Suh, M.C. (2013) Recent advances in cuticular wax biosynthesis and its regulation in Arabidopsis. Mol. Plant, 6, 246–249.

Leide, J., Hildebrandt, U., Reussing, K., Riederer, M. and Vogg, G. (2007) The developmental pattern of tomato fruit wax accumulation and its impact on cuticular transpiration barrier properties: Effects of a deficiency in a *β*-ketoacyl-coenzyme A synthase (LeCER6). Plant Physiol., 144, 1667–1679.

Lewandowska, M., Keyl, A. and Feussner, I. (2020) Wax biosynthesis in response to danger: its regulation upon abiotic and biotic stress. New Phytol., 227, 698–713.

Li, Li, L., Du, Y., et al. (2019) Maize glossy6 is involved in cuticular wax deposition and drought tolerance. J. Exp. Bot., 70, 3089–3099.

Li, L., Li, D., Liu, S., et al. (2013) The maize glossy13 gene, cloned via BSR-Seq and Seq-walking encodes a putative ABC transporter required for the normal accumulation of epicuticular waxes. PLoS ONE, 8, e82333.

Lin, M., Matschi, S., Vasquez, M., et al. (2020) Genome-wide association study for maize leaf cuticular conductance identifies candidate genes involved in the regulation of cuticle development. G3, 10, 1671–1683.

Lin, M., Qiao, P., Matschi, S., et al. (2022) Integrating GWAS and TWAS to elucidate the genetic architecture of maize leaf cuticular conductance. Plant Physiol., 189, 2144–2158.

Lipka, A.E., Gore, M.A., Magallanes-Lundback, M., et al. (2013) Genome-wide association study and pathway-level analysis of tocochromanol levels in maize grain. G3, 3, 1287–1299.

Lipka, A.E., Tian, F., Wang, Q., Peiffer, J., Li, M., Bradbury, P.J., Gore, M.A., Buckler, E.S. and Zhang, Z. (2012) GAPIT: genome association and prediction integrated tool. Bioinformatics, 28, 2397–2399.

Liu, Q., Huang, H., Chen, Y., Yue, Z., Wang, Z., Qu, T., Xu, D., Lü, S. and Hu, H. (2022) Two Arabidopsis MYB-SHAQKYF transcription repressors regulate leaf wax biosynthesis via transcriptional suppression on DEWAX. New Phytol., 236, 2115–2130.

Liu, S., Dietrich, C.R. and Schnable, P.S. (2009) DLA-based strategies for cloning insertion mutants: cloning the gl4 locus of maize using Mu transposon tagged alleles. Genetics, 183, 1215–1225.

Liu, S., Yeh, C.-T., Tang, H.M., Nettleton, D. and Schnable, P.S. (2012) Gene mapping via bulked segregant RNA-Seq (BSR-Seq). PLoS ONE, 7, e36406.

Lubin, J.H., Colt, J.S., Camann, D., Davis, S., Cerhan, J.R., Severson, R.K., Bernstein, L. and Hartge, P. (2004) Epidemiologic evaluation of measurement data in the presence of detection limits. Environ. Health Perspect., 112, 1691–1696.

Lundqvist, U. and Lundqvist, A. (2008) Mutagen specificity in barley for 1580 eceriferum mutants localized to 79 loci. Hereditas, 108, 1–12.

Luo, X., Wang, B., Gao, S., Zhang, F., Terzaghi, W. and Dai, M. (2019) Genome-wide association study dissects the genetic bases of salt tolerance in maize seedlings. J. Integr. Plant Biol., 61, 658–674.

Luo, Z., Tomasi, P., Fahlgren, N. and Abdel-Haleem, H. (2019) Genome-wide association study (GWAS) of leaf cuticular wax components in Camelina sativa identifies genetic loci related to intracellular wax transport. BMC Plant Biol., 19, 187.

Lü, S., Song, T., Kosma, D.K., Parsons, E.P., Rowland, O. and Jenks, M.A. (2009) Arabidopsis CER8 encodes LONG-CHAIN ACYL-COA SYNTHETASE 1 (LACS1) that has overlapping functions with LACS2 in plant wax and cutin synthesis. Plant J., 59, 553–564.

McFarlane, H.E., Shin, J.J.H., Bird, D.A. and Lacey Samuels, A. (2010) *Arabidopsis* ABCG Transporters, which are required for export of diverse cuticular lipids, dimerize in different combinations. Plant Cell, 22, 3066–3075.

McFarlane, H.E., Watanabe, Y., Yang, W., Huang, Y., Ohlrogge, J. and Lacey Samuels, A. (2014) Golgi- and trans-golgi network-mediated vesicle trafficking is required for wax secretion from epidermal cells. Plant Physiol., 164, 1250–1260.

Moose, S.P. and Sisco, P.H. (1994) Glossy15 Controls the Epidermal Juvenile-to-Adult Phase Transition in Maize. Plant Cell, 6, 1343–1355.

Neter, J., Kutner, M.H., Nachtsheim, C.J. and Wasserman, W. (1996) Applied Linear Statistical Models, Irwin Chicago.

Patel, S., Nelson, D.R. and Gibbs, A.G. (2001) Chemical and physical analyses of wax ester properties. J. Insect Sci., 1, 4.

Patwari, P., Salewski, V., Gutbrod, K., et al. (2019) Surface wax esters contribute to drought tolerance in Arabidopsis. Plant J., 98, 727–744.

Pollard, M., Beisson, F., Li, Y. and Ohlrogge, J.B. (2008) Building lipid barriers: biosynthesis of cutin and suberin. Trends Plant Sci., 13, 236–246.

Post-Beittenmiller, D. (1996) Biochemistry and molecular biology of wax production in plants. Annu. Rev. Plant Biol., 47, 405–430.

Qiao, P., Bourgault, R., Mohammadi, M., Matschi, S., Philippe, G., Smith, L.G., Gore, M.A., Molina, I. and Scanlon, M.J. (2020) Transcriptomic network analyses shed light on the regulation of cuticle development in maize leaves. Proc. Natl. Acad. Sci. USA, 117, 12464–12471.

R Core Team (2019) R: A language and environment for statistical computing, R Foundation for Statistical Computing.

Resco de Dios, V., Chowdhury, F.I., Granda, E., Yao, Y. and Tissue, D.T. (2019) Assessing the potential functions of nocturnal stomatal conductance in C3 and C4 plants. New Phytol., 223, 1696–1706.

Riederer, M. and Schreiber, L. (2001) Protecting against water loss: analysis of the barrier properties of plant cuticles. J. Exp. Bot., 52, 2023–2032.

Schnable, P.S., Stinard, P.S., Wen, T.-J., Heinen, S. and Weber, D. (1994) The genetics of cuticular wax biosynthesis. Maydica, 39, 279–287.

Schönherr, J. (1976) Water permeability of isolated cuticular membranes: The effect of cuticular waxes on diffusion of water. Planta, 131, 159–164.

Schönherr, J. and Riederer, M. (1989) Foliar penetration and accumulation of organic chemicals in plant cuticles. In G. W. Ware, ed. Reviews of Environmental Contamination and Toxicology. New York, NY: Springer New York, pp. 1–70.

Schuster, A.-C., Burghardt, M. and Riederer, M. (2017) The ecophysiology of leaf cuticular transpiration: are cuticular water permeabilities adapted to ecological conditions? J. Exp. Bot., 68, 5271–5279.

Schwarz, G. (1978) Estimating the dimension of a model. Ann. Stat., 6, 461–464.

Serrano, M., Coluccia, F., Torres, M., L’Haridon, F. and Métraux, J.-P. (2014) The cuticle and plant defense to pathogens. Front. Plant Sci., 5, 274.

Seufert, P., Staiger, S., Arand, K., Bueno, A., Burghardt, M. and Riederer, M. (2021) Building a barrier: The influence of different wax fractions on the water transpiration barrier of leaf cuticles. Front. Plant Sci., 12, 766602.

Stegle, O., Parts, L., Durbin, R. and Winn, J. (2010) A Bayesian framework to account for complex non-genetic factors in gene expression levels greatly increases power in eQTL studies. PLoS Comput. Biol., 6, e1000770.

Stenmark, H. (2009) Rab GTPases as coordinators of vesicle traffic. Nat. Rev. Mol. Cell Biol., 10, 513–525.

Strobl, C., Boulesteix, A.-L., Zeileis, A. and Hothorn, T. (2007) Bias in random forest variable importance measures: illustrations, sources and a solution. BMC Bioinformatics, 8, 25.

Sturaro, M., Hartings, H., Schmelzer, E., Velasco, R., Salamini, F. and Motto, M. (2005) Cloning and characterization of GLOSSY1, a maize gene involved in cuticle membrane and wax production. Plant Physiol., 138, 478–489.

Sun, G., Xia, M., Li, J., et al. (2022) The maize single-nucleus transcriptome comprehensively describes signaling networks governing movement and development of grass stomata. Plant Cell, 34, 1890–1911.

Sun, G., Zhu, C., Kramer, M.H., Yang, S.-S., Song, W., Piepho, H.-P. and Yu, J. (2010) Variation explained in mixed-model association mapping. Heredity, 105, 333–340.

Tacke, E., Korfhage, C., Michel, D., Maddaloni, M., Motto, M., Lanzini, S., Salamini, F. and Döring, H.P. (1995) Transposon tagging of the maize Glossy2 locus with the transposable element En/Spm. Plant J., 8, 907–917.

Taiz, L. and Zeiger, E. (2010) *Plant Physiology*, Sunderland, Massachusetts: Sinauer Associates, Inc.

Tang, J., Yang, X., Xiao, C., Li, J., Chen, Y., Li, R., Li, S., Lü, S. and Hu, H. (2020) GDSL lipase occluded stomatal pore 1 is required for wax biosynthesis and stomatal cuticular ledge formation. New Phytol., 228, 1880–1896.

Thimmappa, R., Geisler, K., Louveau, T., O’Maille, P. and Osbourn, A. (2014) Triterpene biosynthesis in plants. Annu. Rev. Plant Biol., 65, 225–257.

Vogg, G., Fischer, S., Leide, J., Emmanuel, E., Jetter, R., Levy, A.A. and Riederer, M. (2004) Tomato fruit cuticular waxes and their effects on transpiration barrier properties: functional characterization of a mutant deficient in a very-long-chain fatty acid β-ketoacyl-CoA synthase. J. Exp. Bot., 55, 1401–1410.

Winter, V. and Hauser, M.-T. (2006) Exploring the ESCRTing machinery in eukaryotes. Trends Plant Sci., 11, 115–123.

Wu, D., Tanaka, R., Li, X., et al. (2021) High-resolution genome-wide association study pinpoints metal transporter and chelator genes involved in the genetic control of element levels in maize grain. G3, 11, jkab059.

Xu, L., Hao, J., Lv, M., et al. (2024) A genome-wide association study identifies genes associated with cuticular wax metabolism in maize. *Plant Physiol.*, kiae007.

Xu, X., Dietrich, C.R., Lessire, R., Nikolau, B.J. and Schnable, P.S. (2002) The endoplasmic reticulum-associated maize GL8 protein is a component of the acyl-coenzyme A elongase involved in the production of cuticular waxes. Plant Physiol., 128, 924–934.

Yang, X., Cui, L., Li, S., Ma, C., Kosma, D.K., Zhao, H. and Lü, S. (2022) Fatty alcohol oxidase 3 (FAO3) and FAO4b connect the alcohol- and alkane-forming pathways in Arabidopsis stem wax biosynthesis. J. Exp. Bot., 73, 3018–3029.

Yeats, T.H. and Rose, J.K.C. (2013) The formation and function of plant cuticles. Plant Physiol., 163, 5–20.

Yuan, X., Zhang, S., Sun, M., Liu, S., Qi, B. and Li, X. (2013) Putative DHHC-cysteine-rich domain S-acyltransferase in plants. PLoS ONE, 8, e75985.

Zhang, Z., Ersoz, E., Lai, C.-Q., et al. (2010) Mixed linear model approach adapted for genome-wide association studies. Nat. Genet., 42, 355–360.

Zheng, J., He, C., Qin, Y., et al. (2019) Co-expression analysis aids in the identification of genes in the cuticular wax pathway in maize. Plant J., 97, 530–542.

